# Cardiac toxicity predictions: Safety pharmacologists correlate with the CiPA model

**DOI:** 10.1101/2020.06.11.144238

**Authors:** Hitesh B. Mistry, Jaimit Parikh

**Affiliations:** Division of Pharmacy, University of Manchester, UK; IBM Research Center, New York, USA

## Abstract

There has been a lot of interest and publicity regarding the use of a complex biophysical model within drug development for predicting the TdeP risk of new compounds. Throughout the development of the complex model numerous groups have shown that a simple linear mechanistic model explains the predictive behaviour of complex mechanistic models. That is the input-output relationship is almost linear even when complex kinetic assays are used. We hypothesized that given this linear relationship that scientist would be able to predict the outcome of the biophysical model. The objective of this pilot study was to assess the feasibility of such an analysis but also assess the initial degree of correlation. A set of 15 compounds with diverse ion-channel blocking against 4 ion-channel currents, IKr, ICaL, INa and INaL, was generated. Safety pharmacologists across numerous companies were approached and asked to categorize the TdeP risk of these compounds using only the % block depicted via a bar chart into one of 3 categories: Risk, No-risk or Unsure. 12 scientists participated in the study, of which 11 correlated strongly with the model (11 person ROC AUC range: 0.86-1, 7 scientists had a value >0.9). The combined prediction of all scientists also correlated strongly with the model. These results highlight that the linear input-output relationship can indeed be predicted by the scientist. A future study exploring the degree of correlation with a wider group of scientists and wider set of compounds would be required to get a more precise estimate of the correlation. We hope this initial exploratory study will encourage the community to pursue this idea.

## Introduction

There has been a lot of interest and publicity regarding the use of a complex biophysical model within drug development for predicting the TdeP risk of new compounds[1, 2, 3]. Throughout the development of the complex model numerous groups have shown that a simple linear mechanistic model explains the predictive behavior of complex mechanistic model [4, 5, 6, 7, 8, 9]. That is the input-output relationship is almost linear even when complex kinetic assays are used. Indeed, this has been confirmed as a property of the CiPA in-silico model [10] via a global sensitivity analysis [11]. If many analyses have shown the input-output relationship to be linear [12, 11]; Is a model actually required? We hypothesized that given this linear relationship that scientist would be able to predict the outcome of the biophysical model. Hence, we designed the following initial exploratory study to assess the strength of the correlation between scientist and the CiPA model predictions.

## Methods

A set of 15 compounds with diverse ion-channel blocking against 4 ion-channel currents, IKr, ICaL, INa and INaL, was generated (Figure 1). Safety pharmacologists across numerous companies were approached and asked to categorize the TdeP risk of these compounds using only the % block depicted via a bar chart into one of 3 categories: Risk, No-risk or Unsure. The correlation between Qnet, a metric derived from of the CiPA model response [10], and the scientist’s choice was done in the following way.

**Figure 1:**
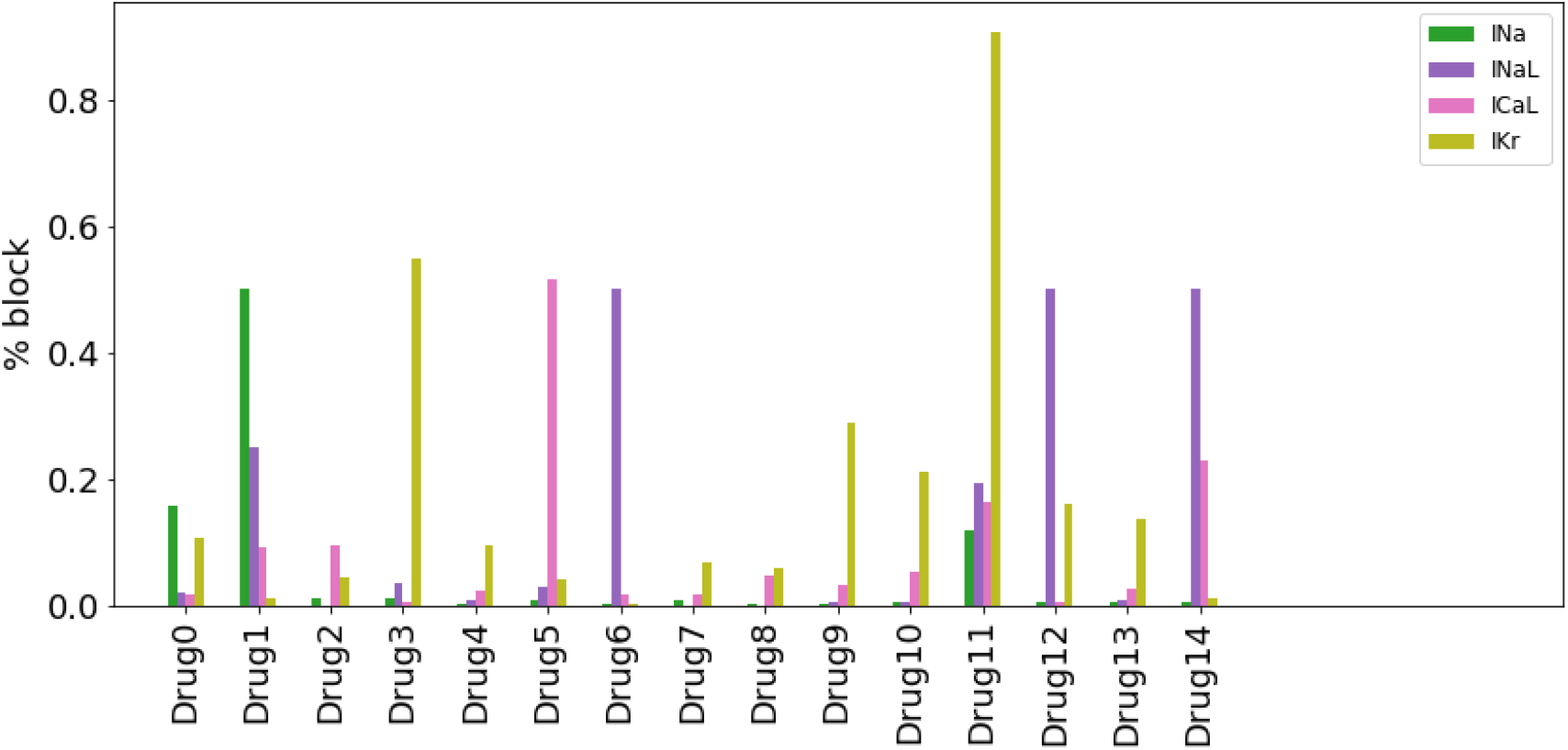
Plot showing the %block of IKr, ICaL, INa and INaL channels for the 15 compounds.

For each scientists’ predictions an ANOVA was conducted with the F-statistic and p-value reported. We also assessed whether Qnet correlated with Risk versus No-Risk/Unsure via the calculation of a ROC AUC reported. In addition, we also grouped all scientists’ predictions by summing up their predictions for each compound, taking the average and comparing this with Qnet values.

## Results

### 15 Compounds

Figure 2 shows the distribution of Qnet across the compound set and the thresholds for the CiPA categorization scheme. The plot shows that the compound set is quite diverse.

**Figure 2:**
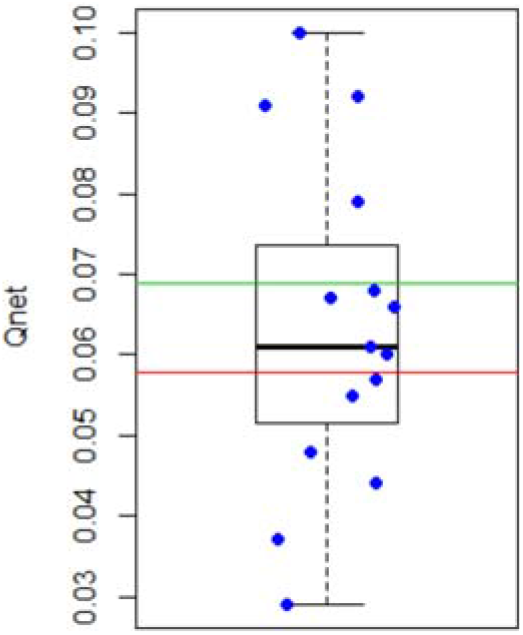
Plot showing the distribution of Qnet values across the compound set. Horizontal lines are the cut-off values from the CiPA validation study.

### Scientist/Model Correlation

Figure 3 shows the response of all scientists across each category for the 15 compounds. Table 1 shows that all scientists except one correlate strongly with Qnet via the two approaches. When we look at consensus across the scientists a strong correlation is found, see Figure 4.

**Table 1:** Individual scientist results

| Scientist | Risk vs<br>Unsure vs No- | Risk vs<br>Unsure/No- |
| --- | --- | --- |

|  | Risk:<br>F-Statistic (P-value) | Risk:<br>ROC AUC |
| --- | --- | --- |
| 1 | 19.5 (<0.001) | 0.98 |
| 2 | 20.8 (<0.001) | 0.96 |
| 3 | 13.3 (0.002) | 0.90 |
| 4 | 20.8 (<0.001) | 0.96 |
| 5 | 12.8 (0.004) | 0.86 |
| 6 | 8.26 (0.013) | 0.89 |
| **7** | **0.02 (0.895)** | **0.50** |
| 8 | 5.33 (0.038) | 0.84 |
| 9 | 22.8 (<0.001) | 1.00 |
| 10 | 10.6 (0.006) | 0.94 |
| 11 | 25 (<0.001) | 0.98 |
| 12 | 10.6 (0.006) | 0.86 |

**Figure 3:**
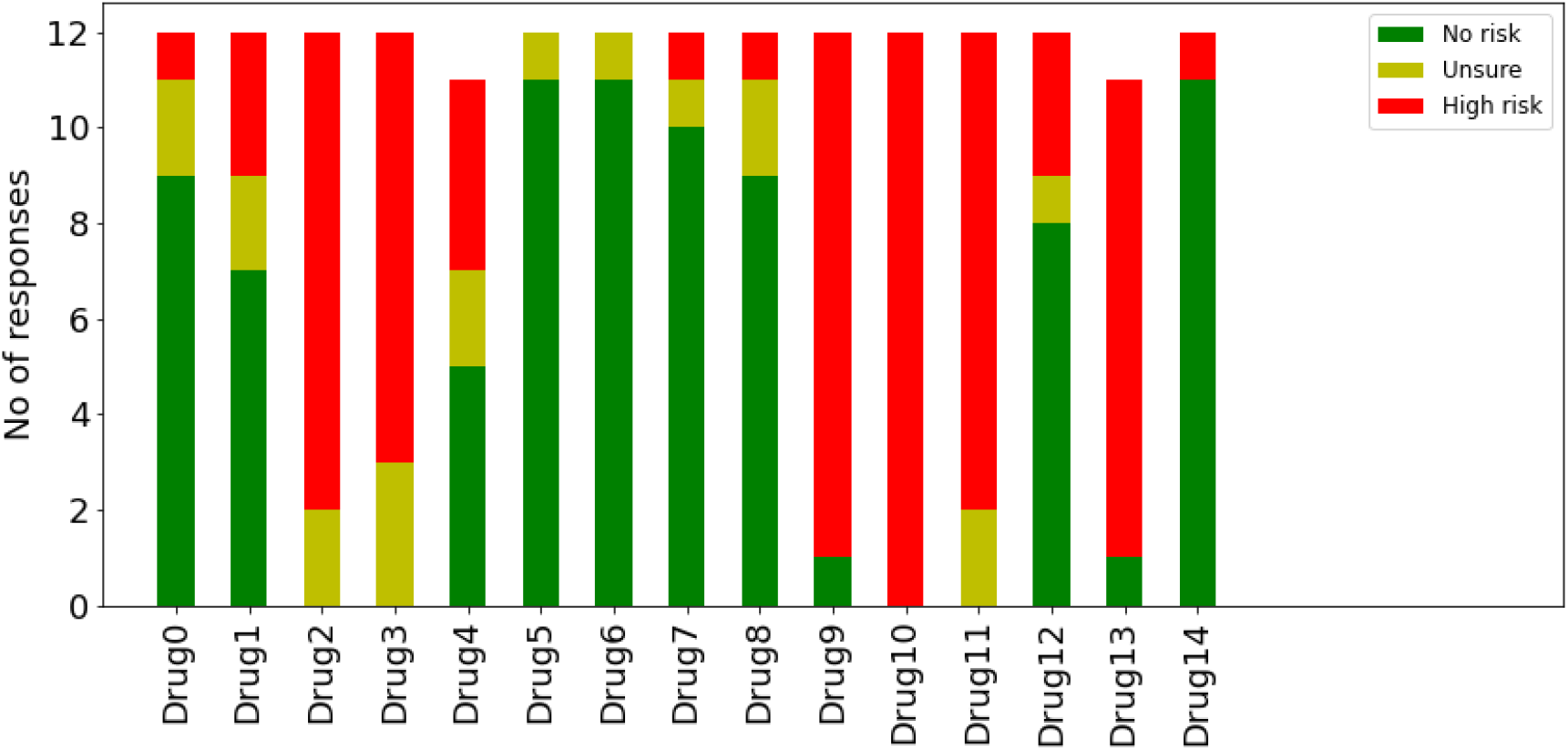
Plot showing the number of responses across different categories by the scientists for each of the 15 drugs.

**Figure 4:**
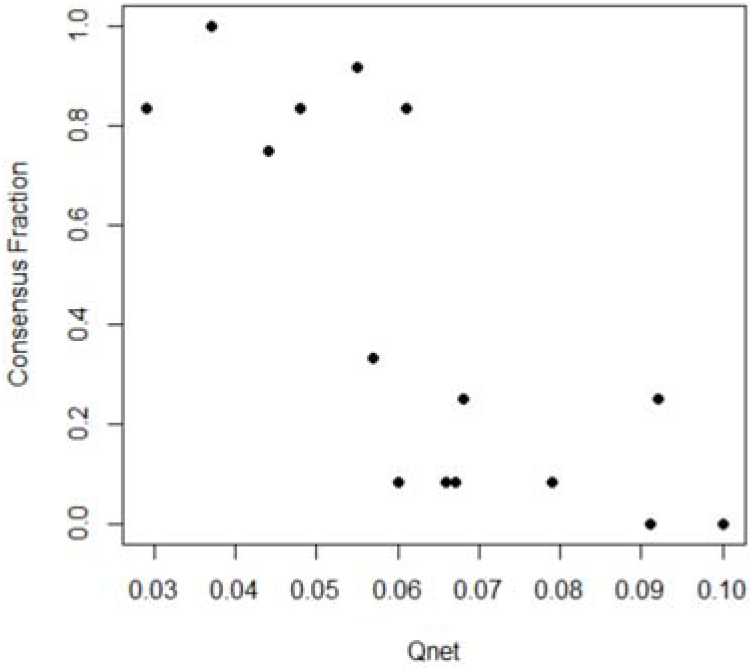
Plot showing the correlation between the Consensus Fraction and Qnet (p=0.001)

## Discussion

Numerous groups have shown that the input-output relationship for the prediction of TdeP using ion-channel pharmacology data is linear [12, 7, 11, 9]. This raises the question as to whether a model is needed at all let alone a complex model. In this initial exploratory study, we have found that scientists predictions do correlate well with Qnet, a complex model metric. This was seen for all scientist questioned except one. We hypothesis that the one scientist whose values did not correlate likely mis-interpreted the question: is there any cardiac risk? However, this may not be the case. One approach to guard against a poor predictor within a group is to take a groups prediction. In our study we found the group prediction via a naïve average did correlate with Qnet.

In summary, these results highlight that the linear input-output relationship can indeed be predicted by the scientist. A future study exploring the degree of correlation with a wider group of scientists and wider set of compounds would be required to get a more precise estimate of the correlation. We hope this initial exploratory study will encourage the community to pursue this idea.

